# Hyphal exploration strategies and habitat modification of an arbuscular mycorrhizal fungus in microengineered soil chips

**DOI:** 10.1101/2023.08.09.552683

**Authors:** Edith C. Hammer, Carlos Arellano-Caicedo, Paola Micaela Mafla-Endara, E. Toby Kiers, Tom Shimizu, Pelle Ohlsson, Kristin Aleklett

## Abstract

Arbuscular mycorrhizal fungi (AMF) are considered ecosystem engineers, however, the exact mechanisms by which they modify and influence their immediate surroundings are largely unknown and difficult to study in soil. In this study, we used microfluidic chips, simulating artificial soil structures, to study foraging strategies and habitat modification of *Rhizophagus irregularis* in symbiotic state associated to carrot roots. Our results suggest that AMF hyphae forage over long distances in void spaces, prefer straight over tortuous passages, anastomose and show strong inducement of branching when encountering obstacles. We observed bi-directional vesicle transport inside active hyphae and documented strategic allocation of biomass within the mycelium e.g., truncated hyphal growth and cytoplasm retraction from inefficient paths. We found *R. irregularis* able to modify pore-spaces in the chips by producing irregularly shaped spores that filled up pores. We suggest that studying AMF hyphal behaviour in spatial settings can explain phenomena reported at bulk scale such as AMF modification of water retention in soils. The use of microfluidic soil chips in AMF research opens up novel opportunities to under very controlled conditions study ecophysiology and interactions of the mycelium with both biotic and abiotic factors.

## Introduction

The small and ancient group of fungi known as arbuscular mycorrhizal fungi (AMF) possess the unique ability to form a mutualistic association with the majority of land plants by trading plant-derived carbon for mineral nutrients (mostly phosphorous and nitrogen) foraged in the soil (Smith and Read, 2008). As a result, they must forage for themselves but also collect resources for their symbiotic partner, and transport both sugar and mineral nutrients through the soil. The plants profit from the fungi’s ability to forage for nutrients that the plant roots cannot access themselves, and due to their much thinner hyphae, fungi have access to a manifold larger volume of the soil pore space than the plants (Smith and Read, 2008).

AMF are exceptional within the kingdom of fungi because of their obligate dependence on plants. As a result, they have two principal types of mycelium: the *intraradical mycelium* which is sheltered inside plant roots, and an autonomous, free growing part, the *extraradical mycelium*, in the soil environment. Physiologically, this intimate relationship to the host plant requires the unique bi-directional long-distance transport of nutrients and carbon compounds within the hyphal structures which must operate simultaneously to re-locate nutrients to the plant and move carbon compounds to the hyphal front for continued foraging.

Energy budgets for foraging are different for AMF compared to other soil fungi: Being fuelled with carbon compounds by the plant means that the AMF likely have a richer and more consistent source of energy, allowing them to forage also in carbon-poor soil spaces. As a result, mycorrhizal fungi are freed of the need to degrade energy-rich organic matter, all the while transporting new biomass into the soil pore system, something which may be important for the build-up of soil carbon (Hawkins et al. 2023 In press). However, because their hyphae must be physically linked to host plants, disturbance and hyphal injury could be even more detrimental for AMF compared to other fungi. The distinct characteristics of AMF and premises under which they forage likely influence the way their hyphae navigate in the soil space and invest their resources. Their different lifestyle has also led to the evolution of an unusual mycelial morphology, with a very sparse distribution of hyphae in the mycelium, a strong hierarchical differentiation of hyphae (ranging from thick runner hyphae of more than 20 µm in diameter down to branched adsorbing structures of less than 1 µm) (Friese and Allen, 1991), and large, oil-filled spores that aid survival rather than spatial dispersal (Smith and Read, 2008; Aguilar-Trigueros *et al*., 2019) though animal and even aerial dispersal also occur (Chaudhary *et al*., 2020).

AMF have been shown to act as ‘ecosystem engineers’ and they can have a profound impact on the physical and chemical makeup of the soil environment: They can modify a soil’s water holding capacity, even beyond their death as necromass (Augé, 2001), its pore space distribution (Bitterlich *et al*., 2018; Pauwels *et al*., 2020), and aggregate soil particles (Wilson *et al*., 2009; Lehmann *et al*., 2017; Rillig and Mummey, 2006). These changes in soil properties are likely caused by interactions between the extraradical mycelium and the surrounding soil matrix, which can have a profound impact on other organisms inhabiting the soil as well as larger biogeochemical processes by e.g. changing water availability (Kakouridis *et al*., 2022). While it is recognised that the symbiotic lifestyle of AMF could have wider effects on the soil ecosystem, and that AMF display different growth- and foraging strategies compared to other fungi, we currently lack a proper understanding of these dynamics. It is therefore of great importance to better understand the ways in which AMF interact with and shape their immediate environment. Studying these processes under realistic conditions has been a major challenge because of the opaque nature of soils, and the fact that their strong spatial complexity and heterogeneity is difficult to mimic under laboratory conditions.

Recent advances in the development of transparent soil micromodels (*soil chips*) enables us to study microbial behaviour and interactions at the scale of their cells (Aleklett *et al*., 2018; Mafla-Endara *et al*., 2021; Schmieder *et al*., 2019; Stanley and van der Heijden, 2017), and to compare mycelial exploration and foraging strategies (Aleklett *et al*., 2021; Held *et al*., 2011, 2019). Here, we used a microengineered soil chip with simulated geometric pore space patterns (Aleklett *et al*., 2021) to study the foraging strategies of *Rhizophagus irregularis* DAOM 197198 at the scale of individual hyphae and follow their interactions within the microstructures. Our aim was to understand basic space exploitation and foraging behaviour of AMF extraradical mycelium. We asked: how do AMF hyphae search and explore uncolonised pore space when foraging in a nutrient void yet complex environment? How do AMF allocate resources within the mycelium during exploration, both in terms of hyphal tip directions and allocation of cytoplasm inside the hyphae? What effects do AMF hyphae have on the spatial pore space in their local environment? To answer these questions we measured (i) the response of individual hyphae to complex microstructures such as narrow channels, sharp turning angles, openings, blockages, and complex obstacle courses, and their interactions with the spatial properties of the pore space (ii) cytoplasm movement and retraction in response to structural confinements at the micro scale, and (iii) the limits of AM fungi exploration distances in an environment that lacks any nutritional reward.

## Methods and materials

### Chip design

To study fungal foraging behaviour, a microfluidic chip called *the Obstacle chip* was designed to test the limitations of exploration and responses to microstructures in fungal hyphae (for a detailed overview of the chip design, see: Aleklett *et al*., 2021; Mafla-Endara *et al*., 2021). Some general features of the chip were that (i) the height of all channels and spaces inside the chip were 7.5µm to avoid overlaying hyphae, (ii) the fungal hyphae always entered through an ‘entry system’ containing round pillars 100 µm wide, spaced 100 µm apart in a grid pattern. This entry system allows fungal hyphae to enter and proliferate but excludes the thicker roots of the root organ culture. Once the hyphae have grown through the entrance system, the chip presents them with a combination of differently shaped channels and obstacles organised in 5 different experimental sections. The experimental sections were; (a) Patches of hexagonal pillars with four different diameters and channel space between them (n=24) (b) Straight channels of 6 different widths (20 µm,15 µm, 10 µm, 8 µm, 6 µm and 4 µm) and all 18.15 mm long, lined with built in rulers to measure growth speed of the advancing hyphae. Each chip contained 5 sets of replicates with the 6 different widths. (c) Channels in patterns with different turning angles, to examine the effect of forced shifts in growth direction. Three types of channels were included: *zigzag* (90° corners, diverting 45° from the original growth direction), *meandering square* (90° corners, diverting 0° or 90° from the original growth direction) and *z-shaped channels* (135° corners, diverting 0° or 135° from the original growth direction and forcing the fungi to repeatedly turn back towards the hyphal front in order to advance) (Fig. 2). Each type of channel was replicated 11 times inside the chip in a randomised order, and with rulers placed between them in order to measure how far the hyphae had advanced. (d) Channels that repeatedly shifted between 10 µm wide straight channels, and 140 µm wide diamond shaped openings. These were designed to examine if hyphal branching or diverging behaviour was induced when possible or in response to hitting an obstacle. One third (12) of the channels contained openings without obstacles, one third (12) of the channels included a 50 µm wide and 10 µm thick obstacle inside the diamond shaped openings, and one third (12) of the channels had a randomised mix of open and blocked diamonds along the channel. Lastly, in section (e), we included two types of mixed obstacle courses within the chip design, which combined multiple kinds of channel-types and obstacles, at two different scales in order to see whether the hyphae were able to find their way to the other end – despite the challenges we had set them. The final design was rendered in AutoCad 2018 (Autodesk), and randomisations of channels inside the chip were achieved using a custom-script provided by UrbanLISP (http://www.urbanlisp.com).

### Chip fabrication, set-up and inoculation

The microfluidic chips were fabricated using molding (full protocol described in Aleklett et al. 2021) and consisted of a soft PDMS (Polydimethylsiloxane) slabs with imprinted channel systems that were UV-bonded to 55×75 mm microscope slides. Bonded chips were placed (a) into 13cm Petri dishes or (b) custom-made glass-bottom one-well plates for high resolution microscopy, as in (Arellano-Caicedo *et al*., 2023) but with microscope slides (Fig. 1 C). We used 4-weeks-old AMF cultures from half Petri dishes (one side of a two-compartmented plate) of root organ cultures (*Daucus carota*) colonised by *R. irregularis* Schenk and Smith (DAOM 197198; Biosystematics Research Center, Ottawa, Canada) in M-medium (Fortin and Becard 1988). The transformed *D. carota* roots originated from a clone of the DC1 line, transformed with T-DNA from the Ri plasmid of *Agrobacterium rhizogenes* (Bécard and Fortin, 1988), originally established by (St-Arnaud *et al*., 1996). Maintenance was accomplished by propagating AM fungi colonised cultures on a minimal nutrient medium (M-medium; Bécard and Fortin, 1988), including 10 g l1 sucrose, low phosphorus concentration of 35 mM (4.8 mg l1 KH2PO4) and 0.3% PhytagelTM for stabilisation (Sigma Chemical Co., St. Louis, MO, USA).

**Figure1.**
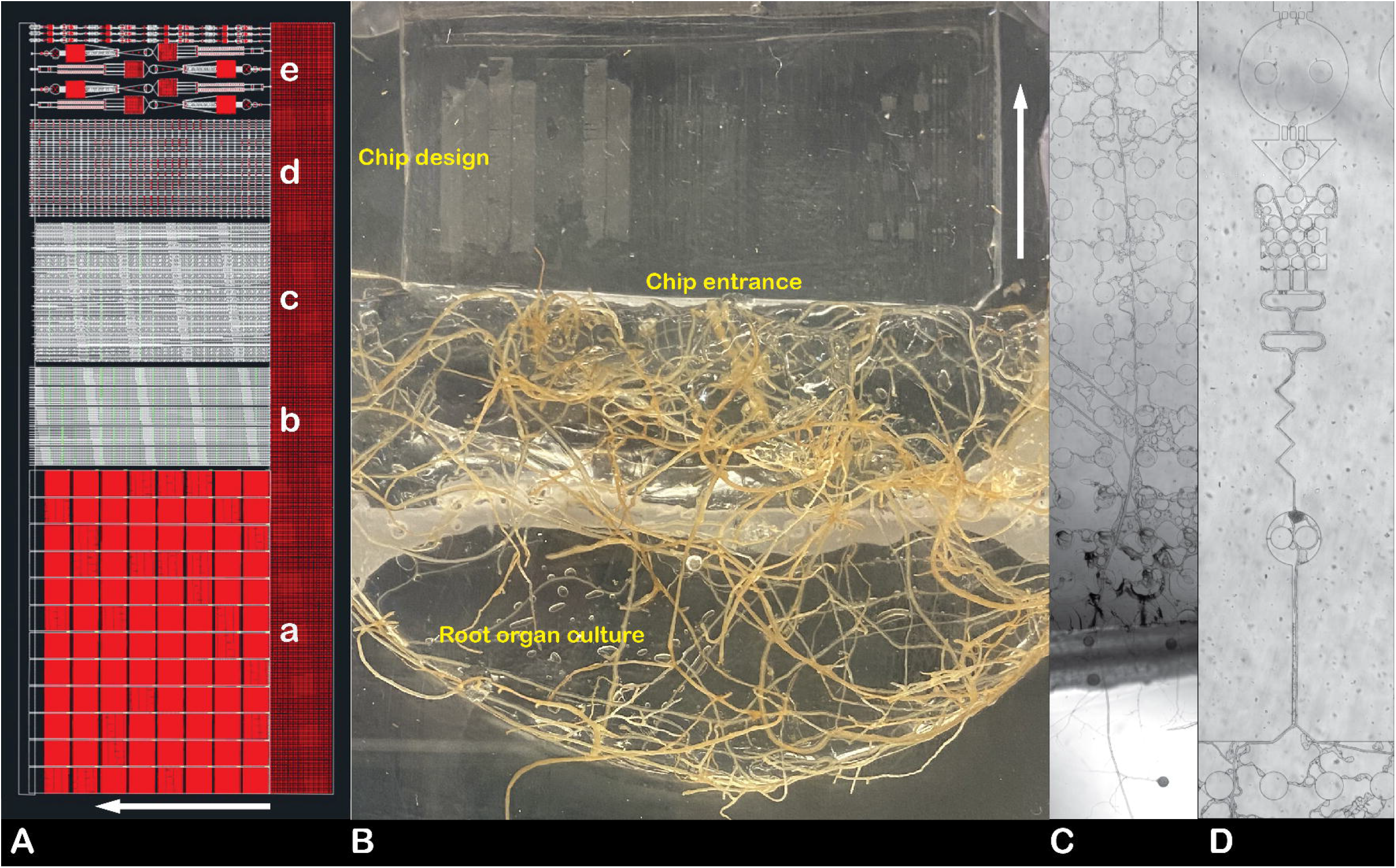
Experimental set-up with *R. irregularis* and the Obstacle chip. (A) Design and layout of the Obstacle chip used in this study, including the different experimental sections a-e (described individually in detail in methods and materials). (B) Microcosm set-up with root organ cultures of *Daucus carota* colonised by *Rhizophagus irregularis* lining the entrance system of the Obstacle chip. In this demonstration, the chip is bonded to a cover slip which is glued into the bottom of a rectangular petri-dish. The petri-dish can then be kept sealed while observations of fungal growth in the chip system are made through the cover slip bottom. (C) Composite image tracing the growth of a hyphae from the phytagel, through the entrance system and further into one of the more complex obstacle courses (D).

Successful colonisation of AMF hyphae into the chip was achieved by placing a half-circle shaped root organ culture with AMF mycelium (cultured in 2-compartmented Petri dishes) next to the opening alongside the chip’s entry system (Fig. 1). Petri dishes containing the chips and the cultures were then sealed with Parafilm to retain moisture within the microcosms.

The narrow height (7.5 µm) of our device limited growth of the mycelium to one focal plane, facilitating microscopic documentation. No water or growth medium was injected into the channels of the chips, hence the only source of nutrients within the system was what the hyphae themselves brought with them or could transport from the roots growing in the initial inoculation plug medium, to study hyphal exploration of an environment without the indication of reward. Root organ cultures adjacent to the chips were supplied with 100 µl sugar solution (4 mg/ml) bi-weekly, and a wet sterile cloth inside the Petri dishes helped to maintain moisture. Microcosms were kept in a dark incubation chamber at 24 °C for optimal growth conditions for the duration of the study.

### Measurements and statistics

Structures in the chips were replicated internally. The designed numbers of replicates in the different experimental sections were: (a): n=24; (b) n=5; (c) n=11; (d) n=36 channels * 32 openings; (e) n=2+2 (two complex obstacle courses and two with flipped start/end sides). Out of these, section b, c and d were systematically examined and used to calculate results in Fig. 2 (n=2 chips) and Fig. 3 (n=3 chips). Sections a and c were only observed to document additional behaviours and sporulation.

**Figure 2:**
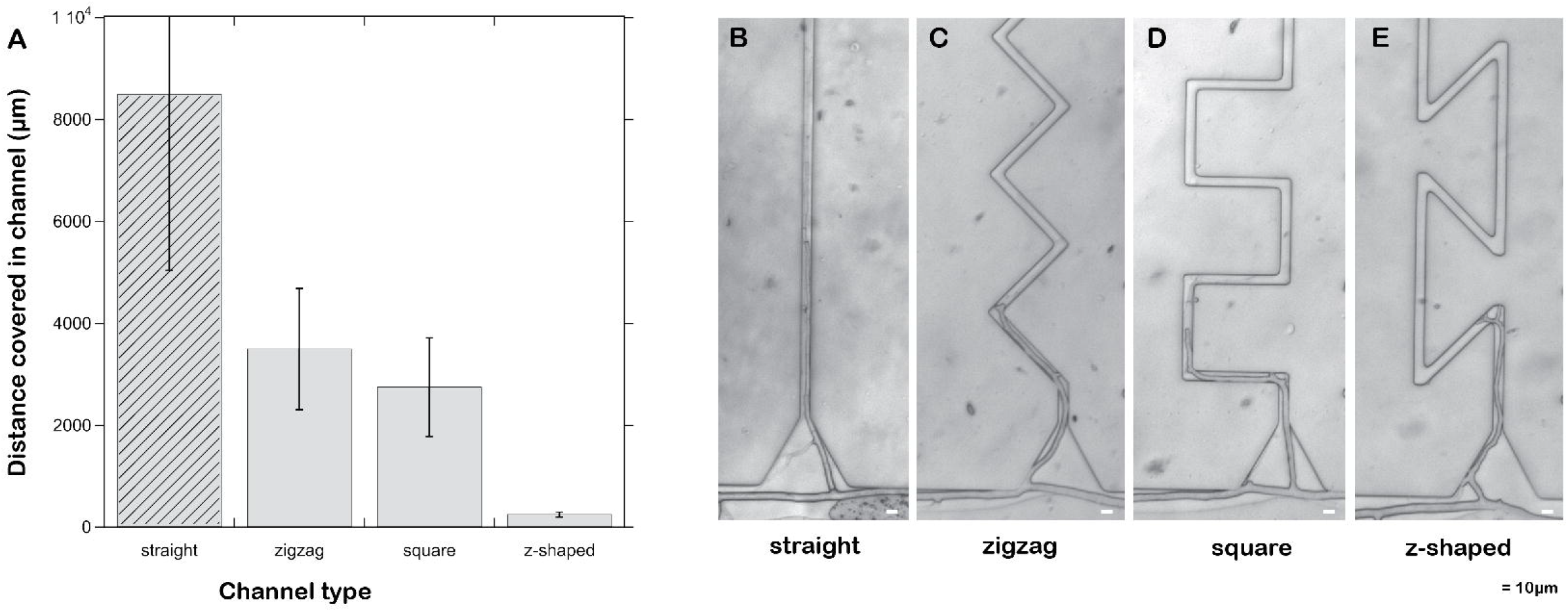
Hyphal growth in 10 µm wide channels with different turning angles. (A) Diagram showing the variation of distances colonised by *R. irregularis* in the various channel-types. We included colonisation results from 10µm straight channels in section b and for comparison (indicated with the diagonally striped pillar). Experimental section (c) includes: *zigzag* (90° corners, diverting 45° from the original growth direction), *meandering square* (90° corners, diverting 0° or 90° from the original growth direction) and *z-shaped channels* (135° corners, diverting 0° or 135° from the original growth direction and forcing the fungi to repeatedly turn back towards the hyphal front in order to advance). (B-E) Example images of hyphal colonisation in the different channel types.

**Figure 3:**
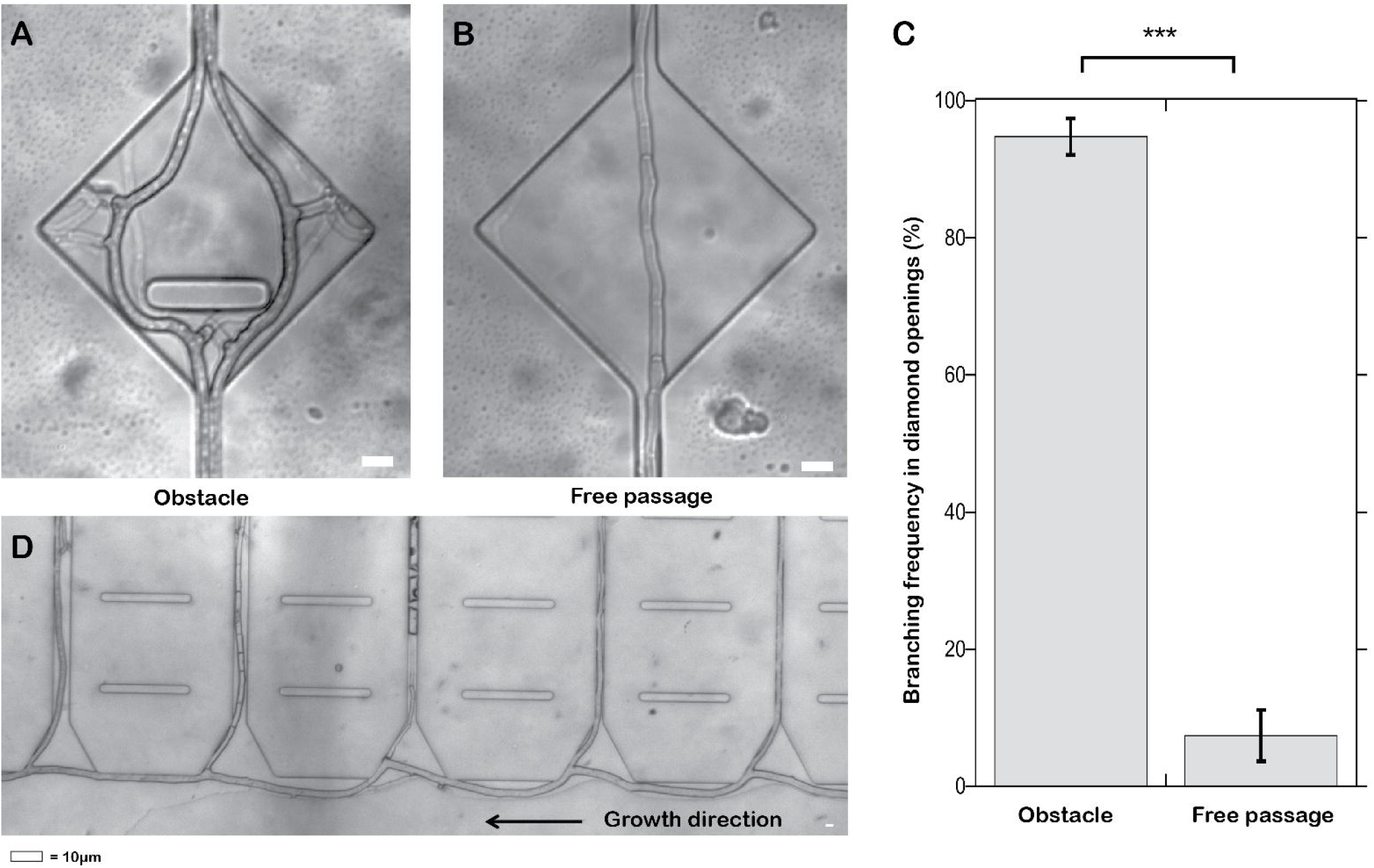
Obstacle induced branching in *R. irregularis*. In experimental section (d), channels repeatedly shifted between being 10 µm wide straight channels, and 140 µm wide diamond shaped openings, some of which included a 50 µm wide and 10 µm thick obstacle. Hyphae growing through the blocked openings showed a strong tendency to branch upon encountering the obstacle (A), whereas hyphae in un-blocked openings generally continued straight through without diverging or branching (B). Overall, when an obstacle prevented straight passage, branching was induced in more than 90% of the cases (C). Hyphae of *R. irregularis* frequently branched when a tip was encountering a physical obstacle in other parts of the chip as well. One example of this was seen when a ‘runner hyphae’ were growing diagonally along channel entries, the hyphal tip would hit the wall of the channel entry and grow side ramifications every time (D).

Data was analysed with ANOVA/ standard least squares models in JMP 16 pro (SAS). In case of unequal variances of the residuals, data was log transformed to meet ANOVA assumptions. Chips were statistically analysed separately as well as in a standard least squares model where internal treatment replication was nested under chip replicate. Under results, statistics for the whole experiments are shown.

### Cytoplasm movement and vesicle tracking

To quantify cytoplasm movement, we tracked movement of the cytoplasm via the movement of lipid-containing vesicles appearing as lighter spots inside the hyphae using video image analysis. 13 videos were recorded in the entrance system of 3-week-old chips using a Nikon Diaphot inverted microscope and a USB29 UXG M BL2.3M black and white camera at 400x magnification and 18.3 frames per second. The videos analysed had a duration ranging between 4 and 46 seconds, and the image processing was done using the free image analysis software (Schindelin *et al*., 2012). Once the videos were imported into FIJI, vesicles inside hyphae that showed the best contrast were selected for tracking. Using the Manual tracking plug-in inside the Tracking tool in the Plug-in menu, between 20 to 30 vesicles were tracked manually in each video. Velocity was calculated for each vesicle during a certain period, depending on how long the vesicle could be followed. To obtain velocity values, the total distance covered was divided by the time spent by the particle to travel that distance. A histogram of the occurrences of measured velocities was plotted for flows both from root to hyphal front and via versa, and a representative video is presented in the results.

## Results

### Colonisation of the Obstacle chip

Extraradical hyphae of *R. irregularis* successfully colonised the internal structures of the obstacle chips and entered the chip entry systems in 66% of the set-up chips. From there, the majority also entered the experimental channels (33% of original setups), enabling us to measure the foraging behaviour of individual hyphae in terms of exploration distances, branching inducement, and morphological differentiations such as sporulation and ramification. We also differentiated between active and emptied hyphae and quantified cytoplasm movements inside the hyphae. Chips were commonly colonised by only a few initial individual tips that over time proliferated as thick, long runner hyphae all along the chips (Fig. 1). From these thick runner hyphae, multiple thinner hyphae branched off sidewards, entering channels and pillar systems (Fig. 3).

### Foraging ranges and speeds

To determine the minimum width that can be colonised by the hyphae, we exposed them to a range of channel widths (4-20 µm; Section b) and found that all channel widths tested could be successfully colonised by *R. irregularis.* While the few hyphae that entered 4 or 6 µm wide channels remained short, with one occurrence of a mean growth distance of 2.4 and a maximum of 5.8 mm, complete colonisation of the chip (18.15 mm) was exclusively found in the wider channels (20 or 15 µm width). However, there was no significant effect of channel width on the maximum distance that the exploratory hyphae covered in this dataset which consists of 2 to 6 colonising (exact numbers in Supplementary material) hyphae per channel width category; least squares regression model nested under chip replicate, p=0.17. It was possible for a few single hyphae to complete the whole chip distance also at other positions in the chip. In five observed cases, hyphae turned and grew back towards the entry in a different channel after they had reached the end. The overall longest recorded hyphae in the chips grown by the same hyphal tip measured a total of 72 mm. Hyphae of AMF can fuse, and in this way create shortcuts or lateral connections within the mycelium (anastomosis). Anastomosis of side branches of hyphae could be regularly observed in the pillar systems (e.g. Fig. 5 B). The longest cytoplasmic connection formed by two anastomosis events inside a chip measured 86 mm, despite the fact that the pore space was void (air filled).

The highest recorded growth rate observed in the chip was 3200 µm per day in a 20 µm wide channel during the initial 5 days after chip inoculation. Individual hyphal tips in the mycelium did not progress synchronously, especially as the cultures grew older. Four weeks after chip inoculation, the fastest hyphae growing in straight channels of different width grew at a speed of 980 µm per day (in an 8µm wide channel), while there was very variable growth in other hyphae of the same mycelium, with a mean growth speed of 33 µm/d, ranging 0 – 310µm/d.

### Cytoplasm movement and retraction

The microfluidic chip systems offered excellent opportunities to study cytoplasm movement live inside the mycelium. The bi-directional movement within single hyphal tubes was frequently observed, and quantified at 4 different regions of the mycelium across the chip. Particle tracking inside hyphae using imaging software showed that vesicle movement was faster in the direction towards the hyphal front (p<0.0001), averaging at 3.7 µm sec^−1^ with the highest velocity measured at 5.3 µm sec^−1^ (Video 1, Fig. 5). By comparison, vesicles moving in the direction towards the host plant moved with an average speed of 2.6 µm sec^−1^ (Fig. 5). Additionally, we observed that vesicles did not move in a continuous flow but were transported inside the hyphae in a more pulsating manner (Video 1).

Retraction of the cytoplasm from apparently inefficient pathways such as anastomosed detours could be observed on multiple occasions and abandoned hyphae were observed to instead contain frequent septation (Fig. 4, Fig. 6 A). Runner hyphae occasionally explored the chip sidewards and produced lateral branchings into channels (Fig 3 D). The hierarchy of the tip of the runner hypha was retained, side-ramifications commonly remained short, and their cytoplasm was retracted within days after entering a channel. In the complex obstacle course structures (Fig 4), we observed that when several paths through a structure were possible, generally only the shortest connection was maintained as cytoplasm-filled, while the other connections were abandoned (Video 2, Fig. 6 B).

**Figure 4:**
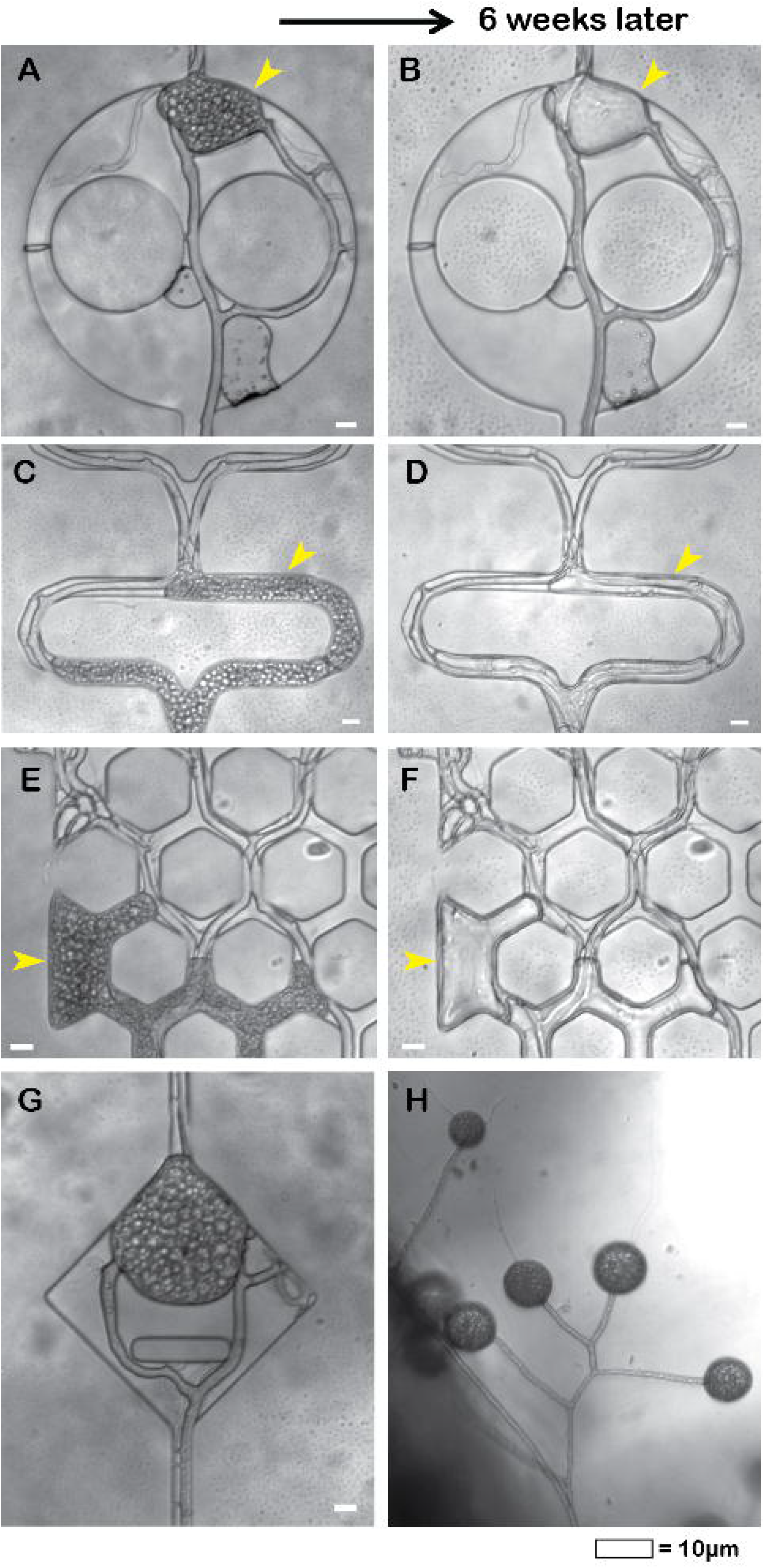
Spore formation of *R. irregularis* inside the obstacle chip. (A, C, E) Pictures showing spores (yellow arrow) formed inside the Obstacle chip in experimental section (e), showing details of the small complex obstacle course and illustrating well the way in which the spores would take on and adapt to the shape of their constricted environment. (B, D, F) Six weeks after the first pictures were taken, the same spots in the chip were re-visited and pictures show that the once lipid-filled spores have been evacuated (yellow arrow). (G) Large spores were also formed in other parts of the chip such as the diamond shaped openings of section (d). (H) While spores inside of the chip had been emptied, those that formed in the phytagel of the adjacent root-organ culture remained round and filled at the same time point, and also showed signs of germination.

**Figure 5:**
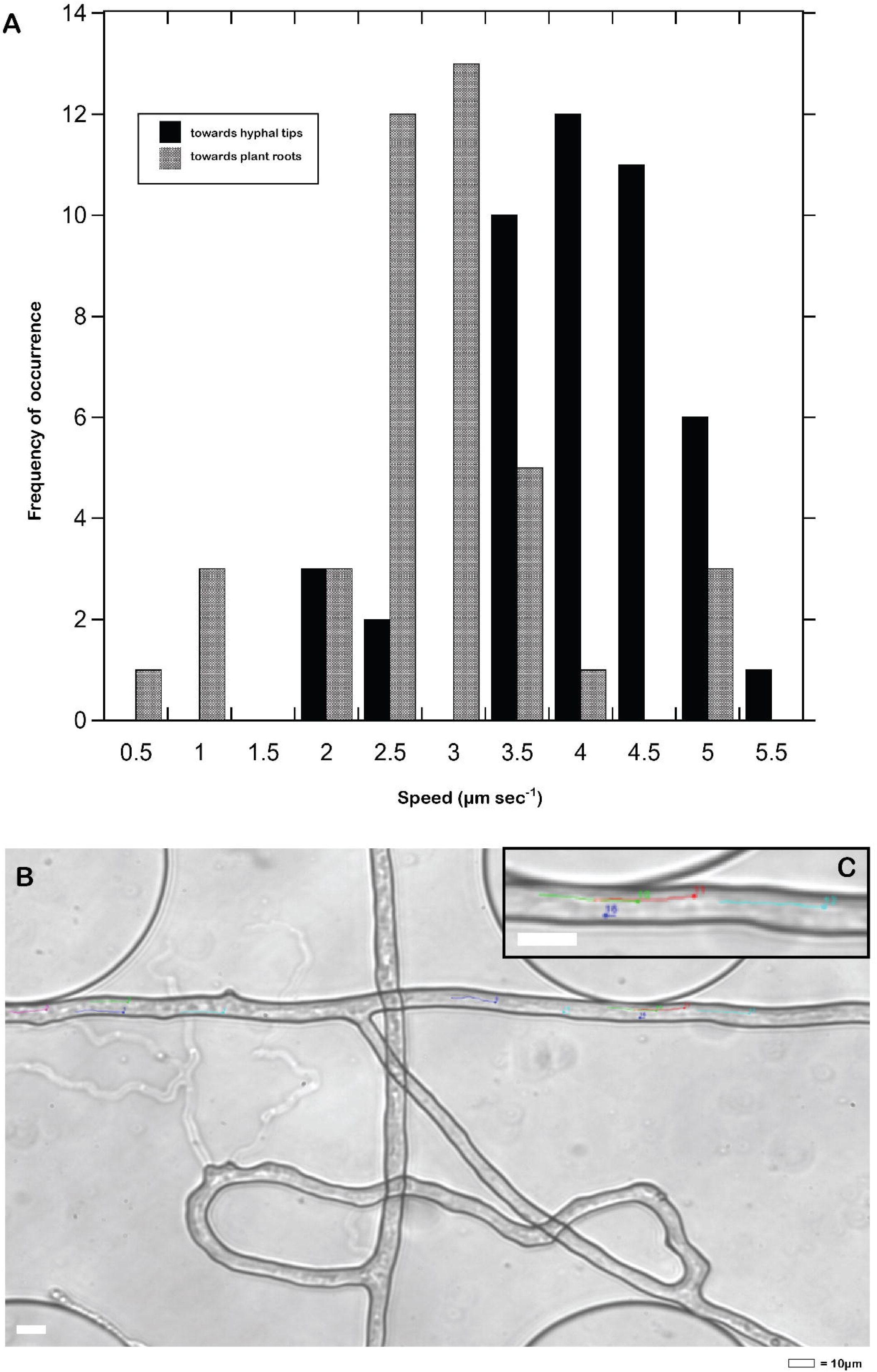
Vesicle movement velocity. (A) Histogram of occurrence frequency of vesicle movement velocities towards the hyphal tip (dark grey) and away from the hyphal tip towards the plant root (light grey). (B) Still frame from Video 1 which was the basis for the vesicle movement analysis and includes examples of two anastomosed hyphae and an abandoned lateral branch. (C) Close-up of identified and tracked vesicles within the hypha.

**Figure 6:**
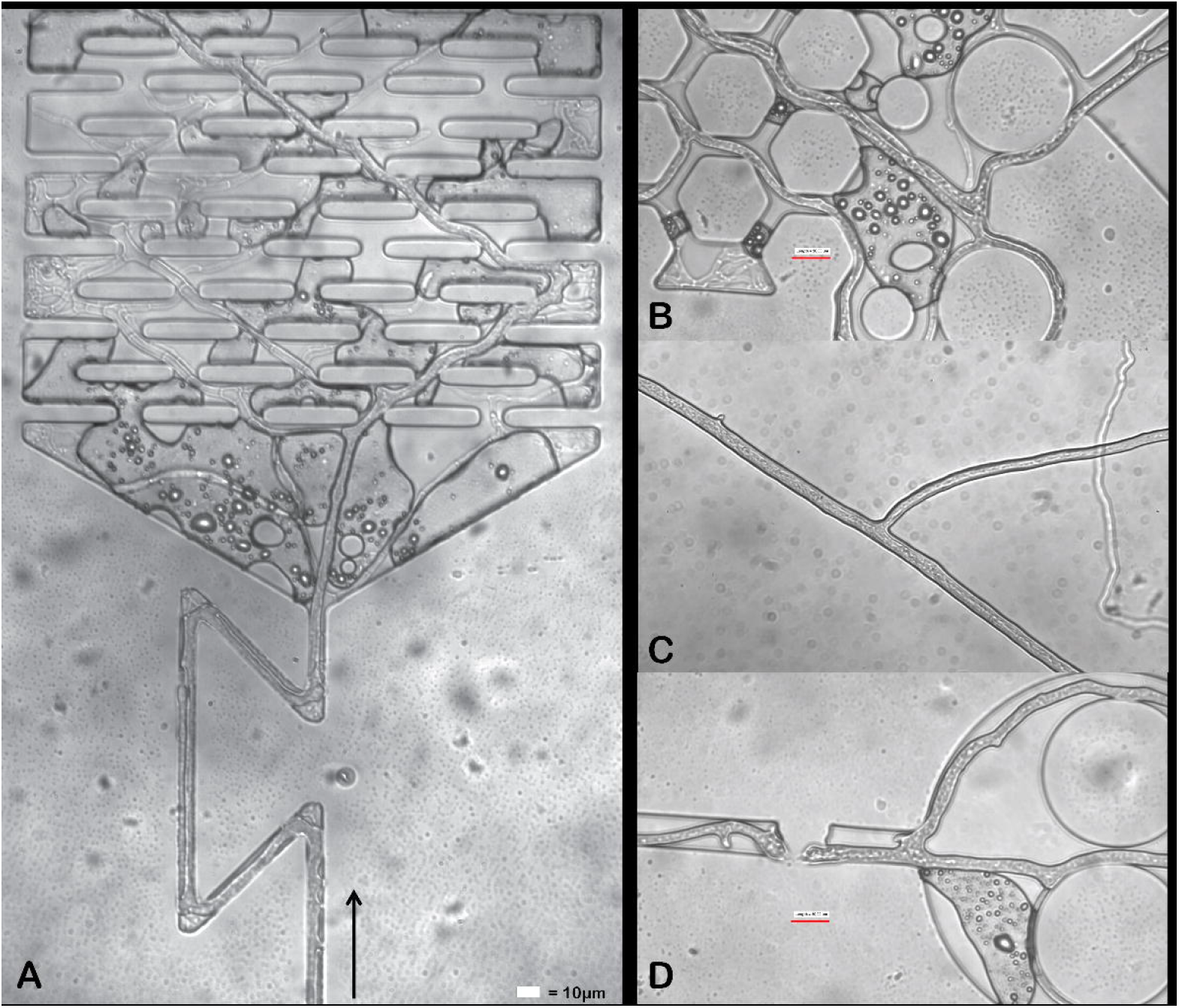
Hyphal colonisation of complex spatial structures and bidirectional cytoplasm movement. Runner hyphae were frequently observed to produce dense branchings in dead-end spaces which were subsequently cut off from cytoplasmic flow through the formation of septae (A, B). (B) Thumbnail of Video 2: Hyphae abandon longer detour paths after anastomosis and occasionally lateral branches. (C) Thumbnail to Video 3: Bi-lateral cytoplasmic stream splits at a lateral branch. (D) Thumbnail to Video 4: Active cytoplasmic transport across a narrow constriction across a fabrication defect of the chip.

### Growth responses to sharp turning angles and corners

We next tested how hyphal growth was affected by growing through channels with corners of different turning angles that in some cases (z-shaped channels) forced them to grow back towards their direction of origin (Section c). We found that in the channels where the AMF hyphae were forced to change their direction of growth, there was a clear preference for growth in square- and zigzag-channels over z-shaped channels (p=0.0099; n=3 chips with 1-6 occurrences per channel category each (see Supplementary material); the distances compared are the actual channel lengths inside the different channel types, not just distance into the chip). Z-shaped channels were often abandoned after the first one or two bends while hyphae reached several mm of distance into the zigzag or square channels (Fig. 2). Hyphae were also commonly observed to produce branchings when they encountered corners. When comparing maximal distances covered in angled channels to the distances covered in the straight channels from Section A, we observed that the distortion of the bends and corners caused hyphae to decrease their foraging distance to less than half (Fig. 2 A).

### Obstacle-induced branching

To test the factors determining diverging and branching patterns, we designed for the hyphae to grow through channels that repeatedly shifted from 10 µm wide straight channels to 140 µm wide diamond shaped halls, spaced 400µm apart (chip section d). Half of the diamonds additionally contained a 50-µm-wide and 10-µm-thick obstacle blocking straight passage for the hyphae. We found that hyphae did not automatically branch or grow sideways when given the opportunity in the wider diamond-shaped sections, but rather directed a straight passage through the opening in 92 % of cases (Fig. 3 A, B). In contrast, when an obstacle was blocking the free passage through the opening, branching occurred in 95 % of the cases (p<0.0001, n=5 chips) When a hypha became trapped in a small dead end (structures such as corners of a patches with hexagonal pillars), it produced strong ramifications that resembled branched absorbing structures (Bago *et al*., 1998) or intraradical arbuscule structures (Fig. 4 B, arrow).

### Sporulation patterns

In two of the colonised chips, spores were formed in several parts of the chip, despite the low height (7.5 µm) of the chip channels. Spores were formed in the diamond-shaped openings, (Fig. 4 G), and in several other spaces of the chip (e.g. within the more complex obstacle courses of Section e, Fig 4 A, C, E) that did not match the spherical form and size of *R. irregularis* spores seen when formed outside of the chip in the media. In the micro-structured chip environment, we found that spore shape was adapted to fit the shape of the channel space (Fig 4 A, C, E). In those cases, spores filled up the whole available volume of that section of the chip and thus blocked off the passage in which it grew. Six weeks after spore production was recorded, most of the spores in the chip had been emptied, and lipids had been retracted from them (Fig. 4 B, D, F). This did not happen to any of the spores formed outside the chip where the fungus grew together with the root in phytagel, even though a few had germinated by that point (Fig. 4 H).

## Discussion

In this study, we demonstrate for the first time the possibilities of growing arbuscular mycorrhizal fungi in microfluidic chips to study the foraging behaviour of their extraradical mycelia. The chips also allow us to quantify intracellular, bi-directional cytoplasmic transport within the mycelial network as done in (Whiteside *et al*., 2019), here in different microspatial settings, and study how re-allocation of biomass occurs as a result of interactions of the mycelium with the soil-pore-space. We further present evidence to suggest that AMF hyphae can drastically modify their immediate micro-habitat by changing pore-space and connectedness through production of large spores that take on a wide variety of shapes based on the surrounding structural limitations. These results open up discussion of how AMF hyphal behaviour might affect larger scale ecosystem processes in the soil and provides a platform for building new experimental designs to study AMF behaviour.

### Characteristics of AMF foraging behaviour and growth patterns

Our experiments revealed features of the extraradical mycelium of AMF that were previously unknown and enable us to relate their hyphal behaviour to other fungal guilds. Foraging behaviours of *R. irregularis* in our study differed from those of other litter decomposing Basidiomycetes previously described and studied in a similar chip design (Aleklett et al. 2021). Hyphal density was markedly lower for AMF in chips compared to most of the litter decomposers examined, especially compared to species like *Coprinellus angulatus* or *Leucopaxillus gentianeus* which generally advanced in synchronised fronts of hyphal tips. Hierarchical and strongly variable hyphal diameters as seen in *R. irregularis* were only rarely observed in the litter decomposers examined (e.g. *Gymnopus confluens* changing from thick sparsely branching hyphae to slimmer frequently branching hyphae further into the chip). One clear difference in behaviour observed between the AMF and the litter decomposers was that *R. irregularis* consistently branched more frequently when faced with an obstacle (87% higher frequency), but not in response to a simple widening of the channel (offering more space to explore). For the litter decomposing basidiomycetes examined in Aleklett et al. 2021, only 2 out of 5 species showed a higher frequency of branching when faced with an obstacle, and increases were small (9 or 16% higher if presented with an obstacle compared to open pore spaces). The branching triggered by environmental physical cues in AMF may compensate for an otherwise fragmentary space exploration coverage via their sparse mycelial architecture.

The frequent apical branching in *R. irregularis* also led to frequent changes of directionality within the mycelium, but despite this, *R. irregularis,* similarly to the Basidiomycetes, struggled with growing through z-shaped channels, and preferably colonised the 90° zigzag and square channels when given the option (Fig. 2). Some of the branching patterns that arose in the Obstacle chip colonised by *R. irregularis* had striking similarities with the branching structures found in AMF, the extraradical branched absorbing structures and the intraradical arbuscules inside plant cells (Fig. 6 A, B). It would be interesting to investigate further if there could be structural and not only chemical cues acting on arbuscule formation during plant-host colonisation by AMF.

Friese and Allen (1991) studied the growth patterns and hyphal architecture of AMF extraradical mycelium of various AMF species and suggested that there are two major forms of hyphae that grow out into the soil from roots; *Runner hyphae* and *absorptive hyphal networks*. Much like the hyphae we found colonising our chips (Fig.1, Fig 3 D), runner hyphae are described as single 10-15 µm wide hyphal strands with angular projections, growing several centimetres out into the soil to infect new roots or tracking along roots to produce multiple infections (Fries and Allen 1991). By comparison, adsorptive hyphal networks are considered to be a series of dichotomously branching hyphae that grow out from the root to form a fan-shaped network in the soil and then die back 5-7 days after formation (Fries and Allen 1991). What we observed in our chip systems was that the architecture of the runner hyphae could diversify into finer networks of thinner hyphae that explored more intricate and complex channels and pore spaces in the chip but that in most cases eventually got shut off from the cytoplasmic flow by the main runner hyphae and died back (Fig. 6 A, B). We did not see clear examples of adsorptive hyphal networks likely due to the lack of nutrient deposits in the chip.

We show that a single runner hypha has the capacity to explore many centimetres of nutrient free air space while maintaining a cytoplasmic connection to the base of the host root. Previous studies of extraradical exploration distances have demonstrated hyphal transport of phosphorus and zinc to the plant host bridging a 15 cm soil compartment (Jansa et al. 2003), and that AMF hyphae can bridge 3.2 mm wide airgaps, while maintaining connection with and transport of water to the plant host (Kakouridis et al., 2022).

### Cytoplasmic movement in AMF and re-allocation of biomass

Fungi have been termed “amoebae in a tunnel shell” since they can move their cytoplasm through their chitin tunnel walls to those areas where physiological activity is most rewarding, and leave empty hyphal shells in old parts of the mycelium. We were able to observe this phenomenon live in our chips (Fig 2b) and report cytoplasm retraction from sub-optimal pathways such as detours in an anastomosed network, with the simultaneous gradual development of septae along the emptied hyphal shells (Fig. 2, 3). The retraction of cytoplasm from freshly built spores that we observed in the chip is to our knowledge the first report of this kind and opens interesting questions on the regulation of resource storage, mobilisation and resumption.

We observed that vesicle movement was faster in the direction of the hyphal front (Fig. 5, four out of four investigations), where the mycelium is expanding and investing biomaterial. We observed that the flow inside the hyphae was not always steady and uniform but could occur in a pulsating manor (Video 1). Pulsating movement of vesicles inside lipids has also been documented in germ tubes of AMF, with irregular motion being more frequent closer to hyphal tips (Bago *et al*., 2002). Since it is thought that apical wall expansion drives cytoplasmic flow inside hyphae (Ramos-García *et al*., 2009) and pulsating hyphal tip elongation has been documented in a collection of different fungal lineages (López-Franco *et al*., 1994), this could be part of explaining the pulsating movement of vesicles seen inside the hyphae.

### AMF habitat modification

Arbuscular mycorrhizal fungi are known to modify their direct environment in multiple ways, including soil aggregation (Rillig and Mummey, 2006), and modification of the water holding capacity of the soil they are growing in (Augé, 2001; Bitterlich *et al*., 2018). The sporulation we observed in openings of the chip design visibly clogged passages and would have prevented any movement of organisms, water or gas across to the adjacent pore space. Irregularly shaped spores of *R. irregularis* have been documented in soils, including variable globose, oblong and irregular, sometimes even knobby shapes (INVAM, Walker, 2013). While it is recognised that different AMF species produce a wide diversity of spore shapes and sizes (Smith and Read, 2008; Aguilar-Trigueros *et al*., 2019), the potential implications of spores adapting their shape and size to the micro scale environment has not previously been examined. In a natural soil environment, AMF spores taking on the shape of the environment may clog passages in the soil pore space – either within or between aggregates. This could result in the alteration of the pore size distribution, organism dispersal and water holding potential of the soil. Alternatively, this phenomenon could partially be an artifact of the fungus growing in the chip system, potentially providing the fungus with more constricted environments than natural soils where the pore space can be modified by force that hyphal tips can apply. Determining the frequency of misshaped spores is difficult because traditional spore extraction procedures may miss out on extracting this type of spores if they eg. are embedded in soil aggregates, and especially if they are only temporarily formed and subsequently emptied with resources redistributed within the mycelium. The fact that only spores in the constricted chip-spaces were emptied and recycled in contrast to the spores in the phytagel (Fig. 4H) suggests some level of recognition from the fungus.

### Culturing AMF in experimental lab systems

Due to the specific growth-requirements of AMF, past work examining foraging behaviour of AMF has been performed in axenic cultures in Petri dishes, most commonly using root organ cultures from carrots (*D. carota*) (Bécard and Fortin, 1988). Under these conditions, it is possible to isolate the extraradical mycelium from the roots, despite their syntrophy. These systems revolutionised the researcher’s possibilities to experimentally manipulate the AMF’s mycelium in the laboratory, and have led to many important insights into the symbiosis, such as morphology and morphology changes (Olsson *et al*., 2014), P transport through the mycelium (Whiteside *et al*., 2019), the rules of the symbiotic trade (Hammer *et al*., 2011; Kiers *et al*., 2011), nutrient distribution in a common mycorrhizal network (Lekberg *et al*., 2010), host effects on nuclear dynamics in the AMF (Kokkoris *et al*., 2021) and interactions with other microorganisms (Elsen *et al*., 2003). However, in root organ cultures, AMF are grown on a homogenous hydrogel nutrient medium lacking any spatial differentiation. This environment is very different from the natural soil habitat of AMF, where fungi are exposed to a structurally, chemically, and biotically complex environment (Kokkoris *et al*., 2019). While still dependent on the low diversity of root organ cultures available for AMF propagation of both plant and fungal species (Goh *et al*., 2022), combining root organ cultures with soil chips allows us to benefit from the latest developments of microfluidic chip technologies that can approach to mimic the physical-spatial and chemical properties of soil or roots at micrometre scale (Aleklett *et al*., 2018).

### The idea of emergent symbiotic behaviours for mycorrhizal interactions

One important question is whether organisms engaged in symbiotic partnerships exhibit traits that are influenced or even controlled on behalf of their partner. For mycorrhizal fungi, the traditional view has been that fungi act as the ‘extended root system of plants’, a view that assumes fungi are subordinate to the plant, and act as passive accessories. However, there is increasing evidence that fungi have evolved their own strategies that allow them actively control interactions with plants, including resource trade patterns (Whiteside *et al*., 2019; Kiers *et al*., 2011; Hammer *et al*., 2011) and choosing their plant partners (Werner *et al*., 2014; Bennett and Groten, 2022). Similarly to how a recent characterisation of mycorrhizal traits define a section of traits as “symbiotic mycorrhizal traits” (Chaudhary *et al*., 2022), one could argue that there exists symbiotic behavioural traits that only manifest when the symbionts are interacting.

In this study we saw that AMF hyphae show a range of different behaviours in how they navigate and explore space, even in a nutrient-void space and when associated with the same plant partner. For example, producing long runner hyphae in search of a new host, branching into arbuscule-like structures in close confinements and re-distributing cytoplasm within the mycelium based on structural cues in the environment (Fig 6A, B, C). Microfluidic chip systems could provide a platform to study how, if, and under which conditions fungal needs are prioritised over plant needs and whether there are emergent symbiotic behaviours for fungi in mycorrhizal associations with different plant species.

### Limitations and future potential of using soil chips for AMF research

Working with both microfluidics and AMF cultures can prove experimentally challenging: Customised chip design and fabrication requires access to microfabrication facilities including clean room facilities, and chip production can be cumbersome. Inoculation of chips with AMF cultures on chips is difficult because of the mycelium’s sparse growth and the need to keep the root organ cultures active and sterile over a long time span. Commitment can however be rewarded since the chips offer possibilities for studies previously not possible: Our work suggests that microfluidic chip systems can be used successfully to study a range of AMF behaviours and interactions, including: (i) *AMF hyphal responses to different stimuli.* Single hyphae can be exposed to controlled conditions of either positive or negative stimuli to study their growth response and compare between different AMF species. (ii) *Details in the process of fungal host recognition.* In microfluidic chip systems, novel inoculation of a root tip can be observed and monitored over time through microscopy and cells can be extracted. (iii) *Interactions between AMF hyphae and other soil organisms* are of high interest, both antagonistic and mutualistic ones. The hyphosphere microbiome includes a broad variety of bacteria living around, and in some cases even within its (Faghihinia *et al*., 2023). Microfluidic chips give us the opportunity to watch, follow and manipulate these interactions in a way that has not earlier been possible at this micrometre scale. (iv) *The study of common mycorrhizal networks* (CMN). CMNs, in which plant individuals are connected through an extraradical mycelium that have either consecutively colonised different roots or fused with other hyphae (anastomosis), have long been a topic of interest in the mycorrhizal field and lately the premises of its existence and function has been the subject of debate (Karst *et al*., 2023). Microfluidic chip systems could potentially provide new ways of studying this phenomenon live in both AMF and ectomycorrhizal fungi (ECM) while still connected to host plants. Channels in microfluidic chips can be designed to funnel sparse hyphae of mycelia towards each other. By tracking individual hyphae, researchers could control and track the extent of anastomosis. Transport and potential transfer of nutrients between the plants could be visualized using quantum dot technology (Whiteside *et al*., 2019) to better understand the dynamics between plants as well as fungal agency in the system.

## Conclusion

By using microfluidic soil chips, we were able to follow the growth and foraging behaviour of *R. irregularis* over time and space. We tracked single hyphae to determine the extents of hyphal colonisation abilities and preferences and observed growth patterns and sporulation in the wider mycelium. We saw evidence of the unique hyphal architecture found in AMF-species (eg. hierarchical runner hyphae and arbuscule-like branchings). Soil chips offer optimal microscopy conditions for sub-cellular studies of AMF, and environmental control at hyphal scale for studying their behaviour expressed at single hyphal tips.

## Supporting information

Supplementary information: replication numbers

## Acknowledgements

ECH recognizes funding from the Swedish Research Council (VR-621-2014-5912), the Foundation for Strategic Research (Future research leader grant SSF FFL18-0089). and the strategic research environment BECC. ETK was supported by an NWO-VICI (202.012), ETK and TS were supported by HFSP (RGP 0029). KA recognises funding from the Swedish Research Council (VR 2022-03505).

