## Supplementary information: replication numbers for "Hyphal exploration strategies and habitat modification of an arbuscular mycorrhizal fungus in microengineered soil chips"

Observed and measured hyphae (n)

Angled channels

| | zigzag | square | z-shaped | p= |
|---|---|---|---|---|
| Chip replicate 1 | 6 | 5 | 2 | 0.06 |
| Chip replicate 2 | 5 | 3 | 4 | 0.009 |
| Chip replicate 3 | 2 | 1 | 1 | 0.28 |

Straight channels

| Width | 20 | 15 | 10 | 8 | 6 | 4 | p= (Regression) | p= (ANOVA) |
|---|---|---|---|---|---|---|---|---|
| Chip replicate 1 | 5 | 5 | 1 | 1 | 1 | 2 | 0.63 | 0.63 |
| Chip replicate 2 | 1 | 1 | 2 | 2 | 2 | 0 | 0.87 | 0.54 |
